# Muropeptides Stimulate Growth Resumption from Stationary Phase in *Escherichia coli*

**DOI:** 10.1101/611111

**Authors:** Arvi Jõers, Kristiina Vind, Sara B. Hernández, Regina Maruste, Marta Pereira, Age Brauer, Maido Remm, Felipe Cava, Tanel Tenson

## Abstract

When nutrients run out, bacteria enter a dormant metabolic state. This low or undetectable metabolic activity helps bacteria to preserve their scant reserves for future, but also diminishes their ability to trace the environment for new growth-promoting substrates. However, neighboring microbial growth is a sure indicator of favorable environment and thus, can serve as a cue for exiting the dormancy. Here we report that for *Escherichia coli* this cue is the basic peptidoglycan unit (i.e. muropeptide). We show that several forms of muropeptides can stimulate growth resumption of dormant *E. coli* cells, but the sugar – peptide bond is crucial for activity. We also demonstrate that muropeptides from several different species can induce growth resumption of *E. coli* and also *Pseudomonas aeruginosa*. These results, together with the previous identification of muropeptides as germination signal for bacterial spores, makes muropeptides rather universal cue for bacterial growth.

## Introduction

Free-living bacteria can encounter large fluctuations in environmental conditions such as those transitioning from nutrient abundance and scarcity, i.e. feast and famine cycle. When growth substrates are exhausted, bacteria initiate specific developmental programs that prepare them for long period of dormancy. Many Gram-positive bacteria form spores that are very resilient to adverse conditions and can survive hundreds of years ^1^. Gram-negatives’ morphological transition associated with the development of dormant cells is, in general, less drastic, but changes do occur.

Gram-negative bacteria undergo large changes in their gene expression pattern and metabolism when entering stationary phase ^2^. These changes are largely governed by the alarmone (p)ppGpp that changes transcription of many genes, reducing growth-oriented gene expression and increasing survival-oriented one ^3,4,5^. Ribosome synthesis is decreased and later inactive 100S ribosomal particles are formed. DNA replication is also inhibited. At the end of these changes cells are entering dormant state, ready to withstand long period without nutrients.

Much less is known about recovery from dormancy when nutrients become available again. It is now clear that cells display considerable phenotypic heterogeneity in timing of recovery – in clonal population some cells start growing rather quickly while others stay dormant for longer and initiate growth only later ^6,7^. The growth resumption timing has been suggested to rely on stochastic process, but some reports also describe different states of dormancy (shallow and deep dormancy) and suggest that growth resumption from shallow dormancy is quicker ^8,9^. We have shown that the order of cells resuming growth in some conditions is determined by the order they enter stationary phase, indicating a long-term memory effect in *E. coli* ^10^.

Persisters are antibiotic tolerant cells in generally antibiotic sensitive bacterial population ^11^. Consensus is now emerging that persisters are mostly nondividing cells and survive antibiotics due to their inactivity (Balaban et al., manuscript submitted to Nature Reviews Microbiology). They only recover from dormancy and start to grow after a long lag phase and by then antibiotic is usually removed. Given that persisters are held responsible for several recurrent infections ^12,13^ it is important to understand the mechanisms that govern the growth resumption process.

The speed of growth resumption can be influenced by the environment. *E. coli* recovery from stationary phase is quicker in rich medium, leading to fewer antibiotic tolerant cells ^6^. Slow recovery in the presence of non-optimal carbon source can be accelerated by small amount of glucose that probably acts as a signal rather than a nutrient ^10^. In *Micrococcus luteus* growing cells secrete an Rpf protein that can induce growth resumption of dormant cells ^14^ and dormant *Staphylococcus aureus* cells can be resuscitated with the help of spent culture supernatant ^15^. *Bacillus* spores are able to detect nutrients and other molecules through specific receptors and initiate germination in response ^16,17^. Here we describe a growth resumption signal for *E. coli* consisting of muropeptides. These molecules are produced by actively growing cells thereby stimulating growth resumption of cells still in dormancy. Moreover, *E. coli* cells are able to resume growth also in response to muropeptides from other species and so are dormant *Pseudomonas aeruginosa* cells, indicating their role in interspecies signaling. We describe the structural requirements for muropeptide activity and isolate mutants with altered sensitivity to muropeptides.

## Results

### Dividing cells secrete a growth resumption promoting factor

During our studies on growth resumption heterogeneity ^10^ we speculated that growing cells produce a signal that stimulates the growth resumption of still non-growing cells. To directly test this hypothesis we prepared a conditioned medium and tested its effect on cells resuming growth from stationary phase. Stationary phase cells were washed and resuspended in fresh medium. Half of the culture was immediately centrifuged and supernatant was sterilized by filtration to be used as control medium for comparisons. The other half was grown until the middle exponential phase and used for preparation of conditioned medium. This conditioned medium still had enough substrates to support growth and also contained factors secreted by growing cells. We took the cells again from stationary phase and compared their growth resumption in fresh, control and conditioned medium. At first we used our single-cell growth resumption assay ^10^ where cells, carrying two plasmids encoding for fluorescent proteins GFP and Crimson, are grown into stationary phase with Crimson expression induced. After that the Crimson inducer is removed and a fresh carbon source is added together with GFP inducer. Cells that resume growth, manifested by active protein synthesis, initially become GFP-positive and later dilute Crimson by cell division (Figure 1a). GFP-positive cells accumulate clearly quicker in conditioned medium. The same effect is evident when optical density of different cultures is compared – lag phase is shorter in conditioned medium (Figure 1b).

**Figure 1.**
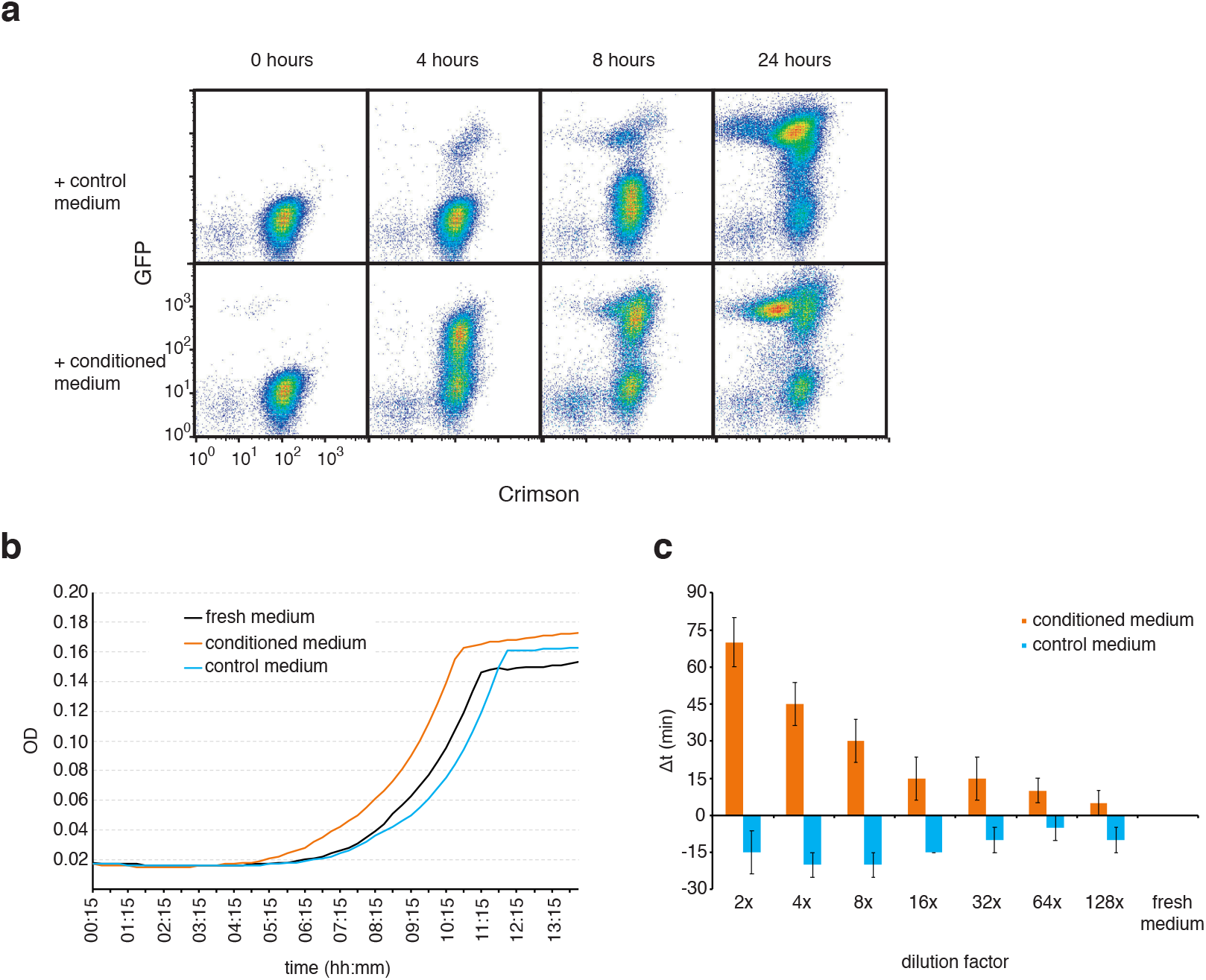
Conditioned medium stimulates growth resumption of stationary phase *E. coli*. **a**. Stationary phase cells were resuspended either in conditioned medium or control medium and GFP expression was induced. Growth resuming cells are GFP-positive and dividing cells have reduced Crimson content due to the dilution by cell division. **b**. Stationary phase cells were diluted into fresh medium, and fresh medium mixed with conditioned medium or control medium. Cells were grown in 96-well plate and OD was measured in every 15 minutes. Cells exposed to conditioned medium resume growth earlier than cells in fresh medium. **c**. Quantification of results in panel b. Δt, the time difference (in minutes) of reaching OD 0.06 (actual reading for 100μl culture) between test medium and fresh medium was calculated. Dilution factor indicates how many times the tested medium was diluted in fresh medium. The average of three independent experiments are shown, error bars indicate standard error of the mean.

Compared to fresh medium, growth resumption is slightly inhibited in control medium. This is probably due to some inhibitory compounds carried over by stationary phase cells (see above how the control medium is made). At the same time the growth rate in exponential phase was the same for all the cultures, indicating that only the growth resumption was affected. For quantification we compared the time it took for different cultures to reach an OD 0.06 (measured OD value for 100 μl culture on 96-well plate, chosen to be approximately in the middle of exponential phase) and calculated the difference between conditioned (or control) medium and the culture grown in fresh medium (Δt). The growth stimulatory effect of conditioned medium was concentration dependent and still clearly detectable when diluted several times (Figure 1c). These results demonstrate that conditioned medium has a growth resumption promoting activity.

### Cell wall derived muropeptides induce growth resumption

We tried to purify this activity from conditioned supernatant, but failed to get enough pure material for identification. However, during this process we learned that the molecule facilitating growth has relatively low molecular weight and is hydrophilic, but not strongly charged. Cell wall derived muropeptides (MPs) fit this description and can induce spore germination in *Bacillus* ^17^, so we tested if MPs can also stimulate growth resumption in *E. coli*. We purified peptidoglycan (PG) from growing *E. coli* cells as described before ^17^ and digested it with mutanolysin to get individual MPs, consisting of a disaccharide (N-acetyl-glucosamine linked to N-acetyl muramic acid) and a short peptide bound to muramic acid (Figure 2a). When added to fresh medium solubilized MPs have a growth resumption promoting activity whereas undigested PG has not (Figure 2b).

**Figure 2.**
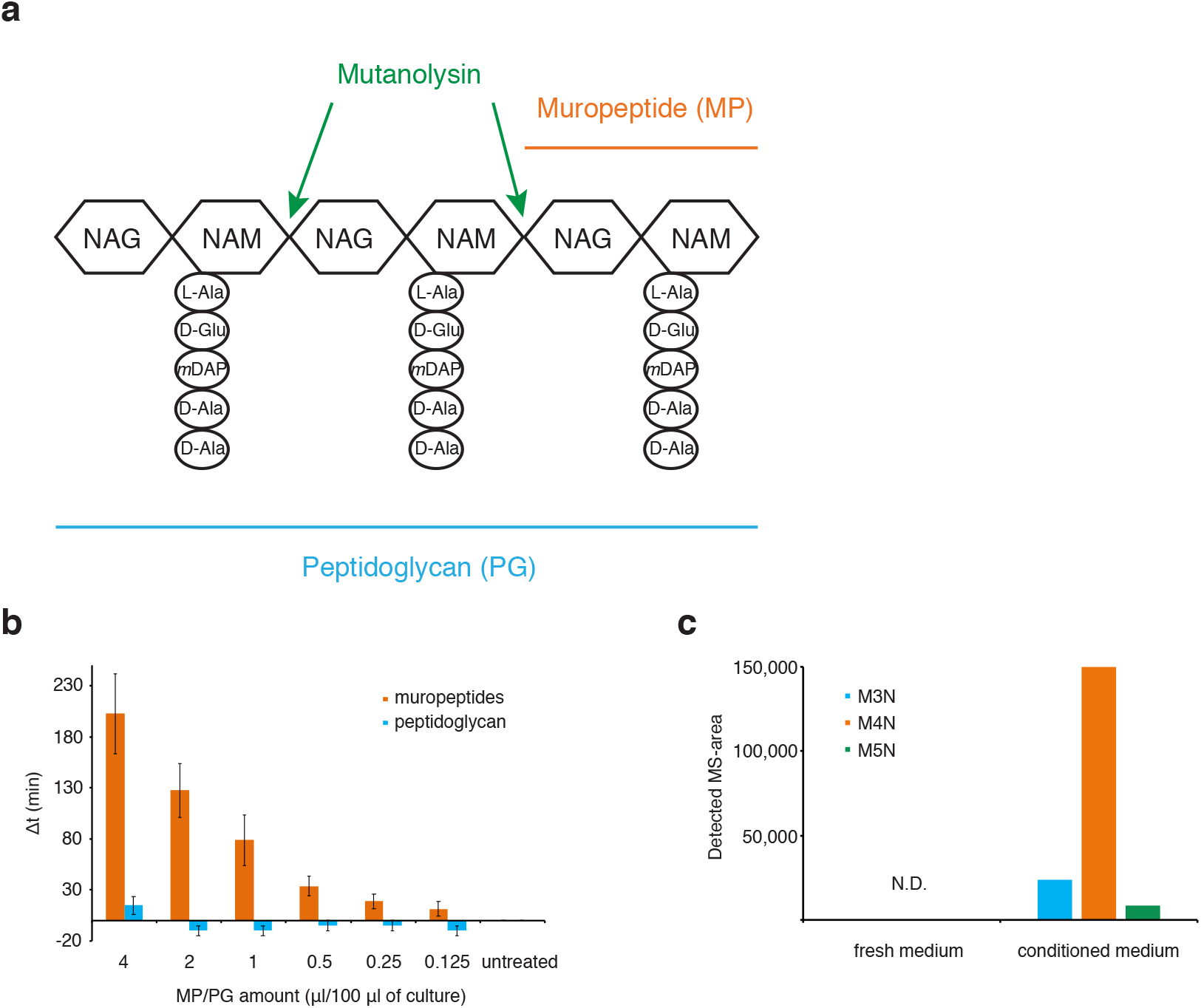
Muropeptides stimulate growth resumption of *E. coli*. **a**. Schematic representation of peptidoglycan and muropeptide. NAG – N-acetyl-glucosamine, NAM – N-acetyl-muramic acid. **b**. Muropeptides (MP) but not peptidoglycan (PG) can stimulate growth resumption of stationary phase cells in fresh medium. Different amounts of MP or PG were added to recovering cells and the Δt was calculated. The average and standard error of the mean of four independent experiments are shown. **c**. Different MP variants can be detected from conditioned medium, but not from fresh medium, using mass-spectrometry. M4N – anhydromuro-tetrapeptide, M3N – anhydromuro-tripeptide, M5N – anhydromuro-pentapeptide, N.D. – not detected. The shown values are an average of two technical replicates.

In order to allow cell enlargement, peptidoglycan hydrolases cleave the PG sacculus during cell growth generating PG fragments. Released MPs are usually recovered – both Gram-positive and Gram-negative bacteria have MP recycling systems that transport PG fragments back to cytoplasm where they can be reused ^18,19^. However, some MPs escape the transport system and are released to growth environment. We analyzed if there are MPs in our conditioned medium. For this, fresh and conditioned medium was concentrated and subjected to UPLC-MS^e^ analysis. We detected anhydro disaccharide with tri-, tetra-, or pentapeptide in conditioned medium, but not in control (fresh) medium (Figure 2c). This is in line with previous results ^20^ and further supports our hypothesis that MPs act as a growth resumption signal in conditioned medium.

It is necessary to notice that the effect of MPs is evident only when the lag phase is long enough. Longer stationary phase, good aeration during the stationary phase and “non-optimal” carbon source in the growth resumption medium all prolong the lag phase and expose the effect of MPs. When stationary phase cells are resuspended in favorable media (LB or MOPS glucose), all cells resume growth quickly and the effect of MPs is not detectable on this background.

Bacterial PG is quite well conserved across different taxa and its basic structure is same in most of the species ^21^. It is thus reasonable to assume that *E. coli* growth resumption could also be stimulated by MPs from other species. One noticeable difference between the PG composition from Gram negatives and some Gram positives is the amino acid present at the third position of the peptide stem: Gram negatives, including *E. coli*, usually have *meso*-diaminopimelic acid (*m*DAP) in that position, while most Gram positives contain L-lysine (Lys) ^21^. We prepared MPs from Gram positive bacteria *Enterococcus faecalis* (contain Lys) and *Bacillus subtilis* (contain *m*DAP) and also from Gram negative *Pseudomonas aeruginosa* (contain *m*DAP). It turned out that soluble MPs, but not PG, from all these species can promote growth resumption of *E. coli* cells (Figure 3a). Furthermore, all these MPs were also able to promote growth resumption of *P. aeruginosa* (Figure 3b). This, together with a fact that conditioned medium from *E. coli* can induce *B. subtilis* spore germination ^17^, indicates that MPs are universal inducers of growth resumption across many bacterial species.

**Figure 3.**
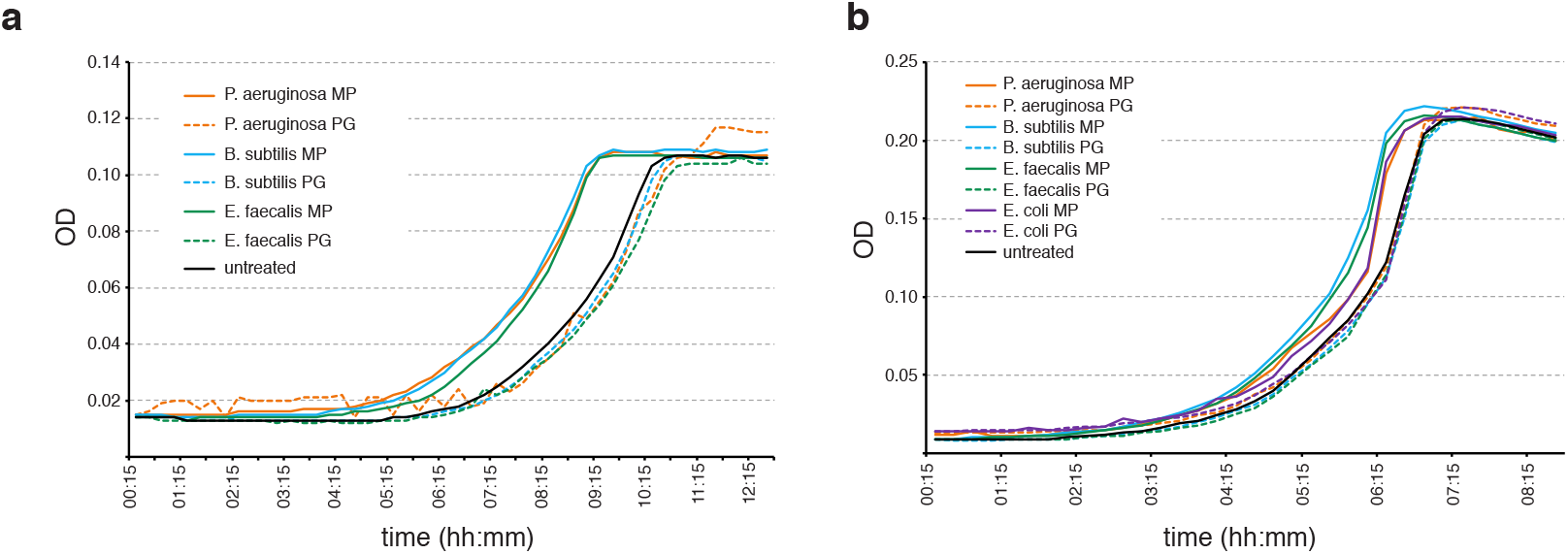
Cross-species recognition of muropeptides as growth resumption stimulators. **a**. Muropeptides from different species stimulate growth resumption of *E. coli* cells. **b**. Muropeptides from different species stimulate growth resumption of *P. aeruginosa* cells. Representative result of at least three independent experiments is shown.

### Sugar-peptide bond of muropeptides is crucial for induction of growth resumption

Digesting PG with mutanolysin results in a mixture of non-crosslinked (monomers) and crosslinked (e.g. dimers and trimers) MPs that can vary in their peptide stem length and composition ^22^. In addition, such a preparation may contain remnants of lipids and proteins associated with PG. To get a better understanding of the activity of different MP variants we purified well-defined structures from the MP mixture and tested their growth resumption properties. Several structures are active in growth resumption assay (Figure 4). Disaccharides with peptides containing 4, 3 or 2 amino acids are all capable of promoting growth resumption and anhydro forms tend to be more active than their hydrogenated counterparts. Even monosaccharide N-acetyl-muramic acid attached to 4 amino acid peptide (anhydro-muramyl-tetrapeptide) can stimulate growth resumption. In contrast, N-acetyl-muramic acid alone or tripeptide alone did not show any growth resumption promoting activity. This indicates that the linkage between sugar and peptide is crucial for the growth resumption activity and if this is present, cells can respond to several MP variants.

**Figure 4.**
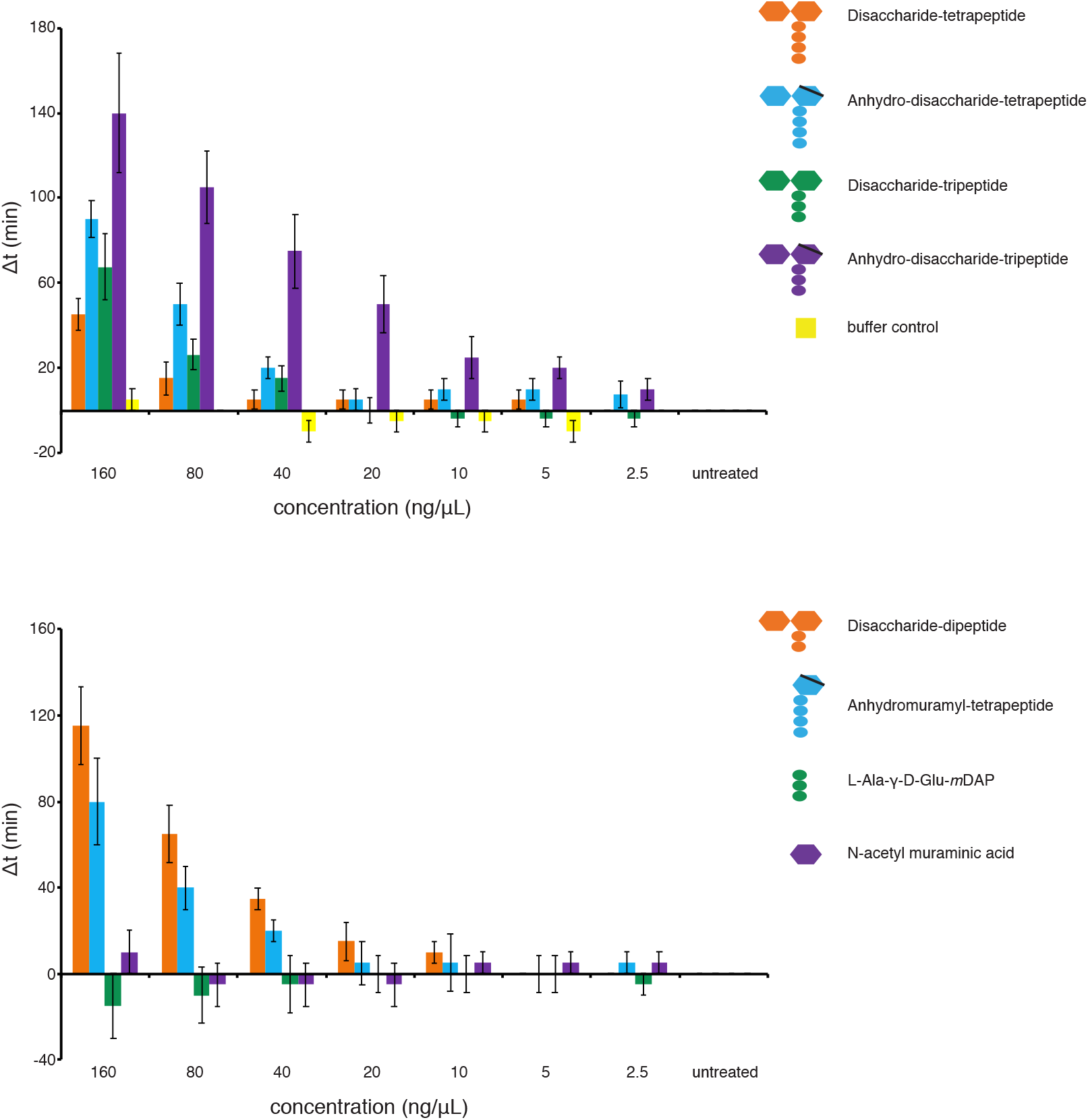
Sugar – peptide bond is crucial for the MP growth resumption activity. Different MP variants were purified and tested on growth resumption assay. Structures with intact sugar-peptide bond can stimulate growth resumption, but NAM or tripeptide alone cannot. The average of three independent biological replicates is shown, error bars indicate standard error of the mean.

### Muropeptide detection by the cells

*E. coli* has a well-described MP recycling system that imports, degrades and recycles anhydromuropeptides released during cell growth ^18^. The genes responsible include *ampG* (permease), *ampD* (amidase), *nagZ* (N-acetylglucosaminidase), *mpaA* (γ-D-Glu-DAP amidase) and *nagB* (glucosamine-6-phosphate deaminase). We expected this system to be responsible also for facilitating MP signaling during growth resumption. However, none of the single knockout strains of genes mentioned above had any phenotype in growth resumption assay and all were responding to MP stimulation like wt (data not shown). This suggests that the primary MP signaling receptor is located either on cell surface or in periplasmic space.

In the case of *Bacillus* spores, MPs are detected through eukaryotic-like protein kinase PrkC ^17^. This gene family is, however, absent in *E. coli*. It is therefore clear that some new pathway for MP detection must be involved in Gram negatives. In order to identify a putative signaling pathway we carried out a genetic screen to find mutants resuming growth relatively slowly in the presence of MPs, but with normal speed in the absence of MPs (see material and methods for details). As a result we identified a clone (BW1.3) whose response to MPs has changed (Figure 5a). BW1.3 cells are more sensitive to MPs at lower concentrations and display a non-monotonous concentration dependency. We sequenced the genome of BW1.3 and identified 2 mutations that result in amino acid changes in two different proteins. In *rpoA* gene, which encodes for RNA polymerase alpha subunit, the amino acid valine was changed to alanine at position 287 (V287A) and in the *yggW* gene, encoding for putative oxidoreductase, the aspartate in position 108 was changed to glutamate (D108E).

**Figure 5.**
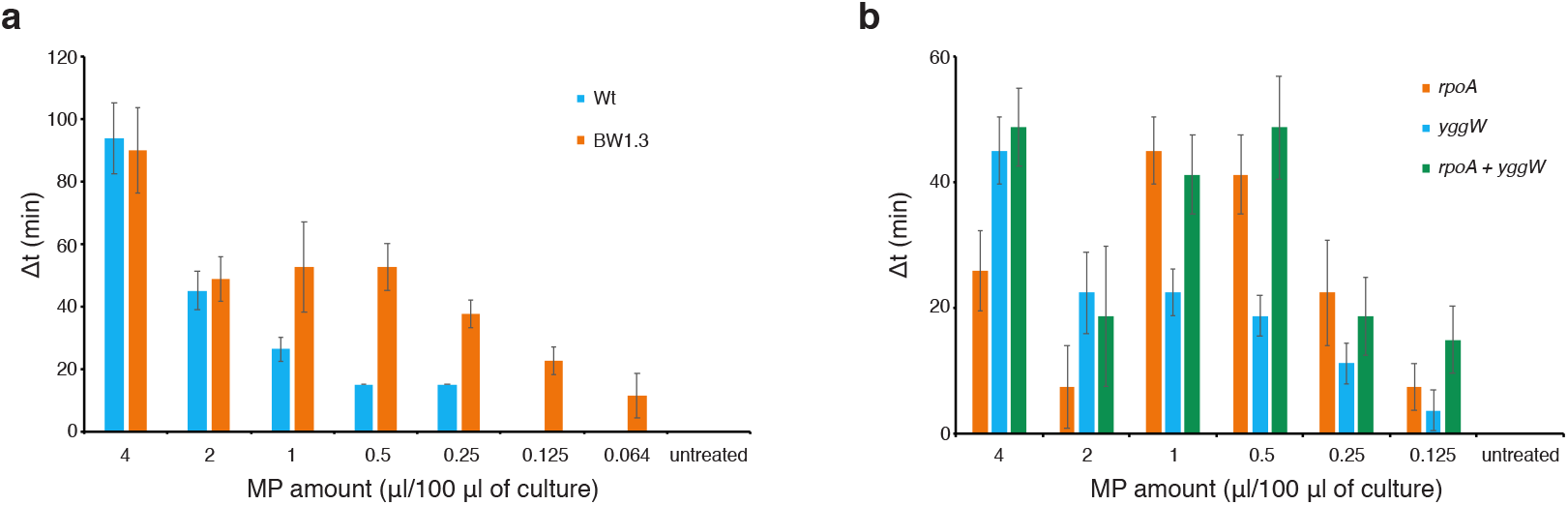
Point mutations in *rpoA* and *yggW* genes alter MP sensitivity. **a**. The strain BW1.3 has an altered sensitivity to MPs. **b**. Point mutations in *rpoA* and *yggW* genes are responsible for change in MP sensitivity. The average of three biological replicates is shown, error bars indicate standard error of the mean.

To validate the role of these mutations we re-introduced these changes to the wt background using CRMAGE technique ^23^. Mutations were made both as single changes and in combination and the resulting strains were tested in growth resumption assay. BW-V287A strain is more sensitive to lower MP concentrations (compared to wt), but the MP effect is dampened at higher concentrations (Figure 5b). BW-D108E is more similar to wt and the double mutant strain combines the two phenotypes displaying a MP effect with dual maxima. V287A and D108E mutations can recapitulate the BW1.3 phenotype and are thus the cause of the change in MP sensitivity.

## Discussion

Dormant bacteria need to monitor their environment in order to detect possible growth substrates. Being ready for all possible ones all the time would be energetically demanding and could deplete the last energy reserves of dormant cells leading to cell death. Growth of other microbes is certainly a sign of favorable environment so detecting others’ growth by some cue could substitute for tracking many possible growth substrates. Here we identify cell wall derived MPs as such cue that can induce growth resumption of dormant cells. We show that in addition to their role as a germination signal for spores ^17^, MPs from both Gram-negative and Gram-positive bacteria can induce growth resumption of *E. coli* and *P. aeruginosa*. The sugar – peptide bond is a crucial structural element for this activity and the detection of MPs by dormant bacteria does not utilize the known MP recycling pathway.

Peptidoglycan hydrolase activity and concurrent solubilization of PG fragments occurs during cell growth ^24^. Both Gram-positive and Gram-negative bacteria have recycling systems that are able to recover some of the released fragments back to the cytoplasm and reuse them ^18,19^. Some of the solubilized products are, however, released to external environment and thus become a cue of cell growth and division. Competing bacteria can also pick up this cue, so it makes sense to curb MP shedding. MP recycling system in *E. coli* is able to work efficiently so that only 6 – 8% of generated MPs are lost at cell division ^20^. Given that PG represents only 2% of cell mass the recycling system does not grant a large energetic advantage and it has been speculated that it has perhaps other specific benefit ^25,18^. Restricting the spread of MPs as a cue for growth-supporting environment could certainly be one.

A scout theory has been proposed as a way for limited population of cells to survive long dormancy while constantly scanning their environment for favorable conditions ^26^. In dormant population some cells randomly exit dormancy and actively scan their environment. If growth is impossible the cell dies after exhausting its energy reserves. However, if the environment supports growth the cell starts to multiply and can signal its siblings to “wake up” and also start growing. MPs certainly fit the role of such signal or cue, although their production seems more unavoidable rather than induced by specific conditions.

MPs act as a germination signal for *Bacillus* spores ^17^. In *Bacillus* they bind to and activate a eukaryotic-like serine-threonine kinase PrkC. This kinase is shown to phosphorylate a two-component system WalRK ^27^, that further controls many genes associated with cell wall metabolism. Mycobacteria have several PrkC homologs, of which PknB is essential for growth ^28^ and can bind PG fragments ^29^. However, PrkC family of proteins is restricted to Gram-positives and *E. coli* or other Gram-negatives do not contain a detectable homolog of PrkC. MP effect in *E. coli* is not facilitated by MP recycling system either, as eliminating the key components of this pathway did not affect the growth resumption effect of MPs (data not shown).

Despite our efforts we were not able to determine a bona fide receptor for MPs in *E. coli*. However, we identified a mutant with altered behavior in response to MPs (Figure 5). A strain carrying single amino acid substitution in the *rpoA* gene (V287A) has a muffled response at the higher MP concentrations, but increased sensitivity at lower concentrations. *rpoA* encodes for RNA polymerase alpha subunit and the region around the position 287 has been implicated in binding to transcriptional activators. Mutating valine 287 to alanine was shown to alter the interaction between RNA Pol alpha subunit and CRP ^30^, increase the FNR-dependent transcription ^31^, and decrease the phage lambda CI-dependent ^32^ and MelR-dependent ^33^ transcription. Correct transcriptional response seems to be necessary for wt-like response to MPs. D108E mutation in *yggW* gene, a putative oxidoreductase, have much milder effect on growth resumption.

The fact that both Gram-negative and Gram-positive bacteria resuscitate in response to PG derived fragments, but detect them through different receptors is an example of convergent evolution. It underlines the importance of keeping track of microbial activity in surroundings by monitoring soluble MPs, as two different detection systems have independently evolved for that function. In addition, eukaryotes have several PG receptors and in mammals NOD1 and NOD2 are used by immune cells to detect the presence of bacteria ^34^. This further emphasizes the importance of MPs as a telltale sign of active bacteria that different organisms have evolved to detect.

## Materials and methods

### Bacterial strains and plasmids

*E. coli* strain BW25113 (F-, Δ(araD-araB)567, ΔlacZ4787(::rrnB-3), λ-, rph-1, Δ(rhaD-rhaB)568, hsdR514) and its derivatives were used in all *E. coli* experiments. Plasmids pET-GFP and pBAD-Crimson ^10^ were used to induce GFP and E2-Crimson expression respectively. In addition, *P. aeruginosa* strain PAO1 was used.

### Growth resumption assay

In the case of *E. coli* cells were grown in MOPS medium supplemented with 0.1% glycerol for 4 – 5 days in 2 mL volume in test tubes. Cells were centrifuged for 1 min at 13,200 rcf, supernatant removed and the pellet was resuspended in equal amount of sterile deionized water. Cells were centrifuged again and resuspended in the same amount of deionized water. Cell suspension was diluted 1:20 in fresh MOPS 0.1% gluconate (or 0.1% glycerol, data not shown) and transferred to 96-well flat bottom plate, 100 μL per well. MPs/PG was added to the first column on the plate and serial dilution was made with two-fold steps. In the case of conditioned medium the first column contained 1:1 mixture of fresh and conditioned medium. Plate was incubated in Biotek SynergyMx plate reader at 37 C degrees with constant shaking. Optical density at 600 nm was measured in every 15 min. during the course of experiment. In the case of *P. aeruginosa* cells were grown in MOPS 0.1% glucose for 5 days and seeded on a 96-well plate in the same medium.

### Flow cytometry analysis

*E. coli* cells with pBAD-Crimson and pET-GFP plasmids were grown in MOPS 0.1% glycerol containing chloramphenicol (25μg/mL), kanamycin (25 μg/mL) and arabinose (1 mM) to induce E2-Crimson. After 4 day incubation in stationary phase cells were washed with water and resuspended either in fresh MOPS 0.1% gluconate or in conditioned medium containing 1 mM IPTG to induce GFP expression. Cells were grown at 37 °C on shaker, samples for flow cytometry were taken at the times indicated, mixed with equal amount of 30% glycerol in PBS and stored at −70 °C pending analysis.

Flow cytometry analysis was carried out as described ^10^ using LSR II (BD Biosciences) with blue (488 nm) and red (638 nm) lasers. The detection windows for GFP and E2-Crimson were 530 ± 15 nm and 660 ± 10 nm respectively. Flow cytometry data was analyzed using FloJo software package. At least 20,000 events were collected for every sample.

### Peptidoglycan isolation

PG was purified as described ^17^. Briefly, cells were grown overnight in LB (*P. aeruginosa, B. subtilis, E. faecalis*) or MOPS 0.2 % glycerol (*E. coli*). 100 ml of bacterial culture was centrifuged, washed twice with deionized water (100 ml and 15 ml) and resuspended in 4 ml of 4% SDS. The suspension was boiled for 30 min, incubated at room temperature overnight and boiled again for 10 min in the next day. SDS-insoluble material was collected by centrifugation at 13,200 rcf for 15 min at room temperature. Pellet was washed four times with deionized water, one time in 12.5 mM Na-phosphate buffer (pH 5.8) and resuspended in 1 ml of the same buffer. The resuspended PG was digested with mutanolysin (Sigma) by adding 1kU of enzyme to 0.7 ml of PG suspension and incubating the mixture at 37 °C overnight with constant shaking. On the next day mutanolysin was inactivated at 80 °C for 20 min.

### Detection of muropeptides in conditioned medium by UPLC-MS

Filtered fresh and conditioned media were dried and resuspended in deionaized water (final samples were 10 times more concentrated than the original medium), boiled for 20 min and centrifuged at 14,000 rpm for 15 min to precipitate proteins and insoluble material before UPLC-MS injection. MS data were obtained by using MS^e^ acquisition mode. These data were processed and a built compound library in UNIFI that contains the structure of several anhydro and non-reduced forms of *m*DAP-type mono and disaccharide peptides was used for the search of MPs. For building the compound library the molecular structure of MPs was obtained by using ChemSketch (www.acdlabs.com). After automatic-compound identification, structure of the matched components was verified by search of corresponding fragment ions and comparison of the mass spectra with MS/MS data previously obtained from standard MPs. The area of the MS-chromatogram obtained for each identified MP was considered as the quantitative value.

UPLC-MS was performed on an UPLC system interfaced with a Xevo G2/XS Q-TOF mass spectrometer (Waters Corp.). Chromatographic separation was achieved using an ACQUITY UPLC-BEH C18 Column (Waters Corp. 2.1 mm × 150 mm; 1.7um particle size) heated at 45 °C. As mobile phases 0.1% formic acid in Milli-Q water (buffer A) and 0.1% formic acid in acetonitrile (buffer B) were used and the gradient of buffer B was set as follows: 0-3 min 5%, 3-6 min 5-6.8%, 6-7.5 min 6.8-9%, 7.5-9 min 9-14%, 9-11 min 14-20%, 11-12 min hold at 20% with a flow rate of 0.175 ml/min; 12-12.10 min 20-90%, 12.1-13.5 min hold at 90%, 13.5-13.6 min 90-2%, 13.6-16 min hold at 2% with a flow rate of 0.3 ml/min; and then 16-18 min hold at 2% with a flow rate of 0.25 ml/min. The QTOF-MS instrument was operated in positive ionization mode using the acquisition mode MS^e^. The parameters set for ESI were: capillary voltage at 3.0 kV, source temperature to 120 °C, desolvation temperature to 350 °C, sample cone voltage to 40 V, cone gas flow 100 L/h and desolvation gas flow 500 L/h. Mass spectra were acquired for molecules eluting only after minute 6 (due to the existence of an abundant background molecule eluting at minute 5 in both fresh and active medium) at a speed of 0.25 s/scan and the scan was in a range of 100–1600 m/z. Data acquisition and processing were performed using UNIFI software package (Waters Corp.).

### Muropeptide production and isolation

Pure MPs were obtained through collection of HPLC (high-performance liquid chromatography) separated MP peaks. Disaccharide-tetrapeptide (M4) and disaccharide-dipeptide (M2) were collected from muramidase-digested sacculi of stationary cell cultures of *Vibrio cholerae* and *Gluconobacter oxydans* grown in LB and YPM (yeast peptone mannitol) medium respectively ^35^. Anhydrodisaccharide-tetrapeptide (M4N) was produced by digesting *V. cholerae* stationary phase sacculi with Slt70 lytic transglycosylase ^35^. For obtaining M3 and M3N tripeptides, M4 and M4N were digested with purified *V. cholerae* L,D-carboxypeptidase LdcV (Hernández et al., in preparation) ^36^. Anhydromuramyl-tetrapeptides (anhNAM-P4) were obtained by digestion of M4N with a purified NagZ homolog of *V. cholerae* (Hernández et al., in preparation)^37,38^. For MP collection, reduced MPs were fractionated by reverse-phase HPLC (Waters Corp.) on an Aeris peptide column (250 × 4.6 mm; 3.6 μm particle size; Phenomenex, USA) using 0.1% of formic acid and 0.1% of formic acid in 40% of acetonitrile as organic solvents in 30 minutes runs ^39,35^. Collected fractions were dried completely and dissolved in water. The identity of individual collected MPs was confirmed by LC-MS analysis. Tripeptide Ala-γ-D-Glu-*m*DAP was purchased from AnaSpec and N-acetyl muramic acid from Sigma-Aldrich.

### Screening for mutants not responding to MP

In order to identify genes involved in MP detection we used *E. coli* strain BW25113 carrying pET-GFP plasmid. Cells were grown into stationary phase in the presence of IPTG to induce GFP expression in MOPS 0.1% glycerol. After four days cells were washed with deionized water and diluted 1:20 into fresh MOPS 0.1% gluconate containing MPs. Growth resumption was monitored by GFP dilution method and nondividing (high GFP content) cells were sorted when they constituted approximately 20% of total population. Sorted cells were pooled and subjected to another round of growth resumption, this time without MP addition and no sorting. In the first round we select cells that resume growth slowly even in the presence of MPs, in the second we select for cells that resume growth with normal speed in the absence of MPs. After four rounds cells were streaked on the agar plate and individual colonies were tested in growth resumption assay. In order to identify mutations behind the phenotype we sequenced the genomes of two clones with altered MP sensitivity (BW1.2 and BW1.3) and two clones with wt-like behavior (BW1.4 and BW1.5) together with wt strain.

### Genome sequencing and bioinformatic analysis

The genomes were sequenced using MiSeq platform (Illumina). Wild-type isolate (BW25113) was assembled with SPAdes (version 3.10.1) ^40^ and used as a reference genome. Sequencing reads from isolates BW1.2, BW1.3, BW1.4 and BW1.5 were mapped to BW25113 using bowtie2 (version 2.0.0-beta7) ^41^. SNPs and small indels for each isolate were called using Samtools (version 1.9) ^42^. Retrieved variations were further filtered to keep only those that were present in BW1.2 and BW1.3 but not in BW1.4 and BW1.5 isolates compared to the wild type. Variations in protein coding areas were verified using Sanger sequencing. Unmapped reads from each isolate BW1.2-BW1.5 were assembled *de novo* to ensure that we were not missing other potential phenotype related sequences that were not presented in wild-type reference assembly.

### Genome modification

Two point mutations identified in selection were re-introduced into wt genome using CRMAGE ^23^. This method combines mutation introduction by oligonucleotide (recombineering) and counterselection against wt using CRISPR-Cas9. The presence of mutations was verified by Sanger sequencing.

## Acknowledgements

This work was supported by Estonian Research Council (grant PRG335), and by the European Regional Development Fund (through the Centre of Excellence in Molecular Cell Engineering). Research in the Cava lab is supported by MIMS, the Knut and Alice Wallenberg Foundation (KAW), the Swedish Research Council and the Kempe Foundation. AB and MR were funded by institutional grant IUT34-11 from the Estonian Ministry of Education and Research and the EU ERDF grant No. 2014-2020.4.01.15-0012 (Estonian Center of Excellence in Genomics and Translational Medicine).

## Author contributions

AJ, FC and TT conceived and designed the study. AJ, KV, RM and MP performed microbiological experiments, SBH analyzed and purified MPs. AB and MR performed genome assembly and bioinformatic analysis. AJ, SBH, FC and TT wrote the manuscript.

